# In-silico studies of Riparin B in the design of drugs: Physicochemical, pharmacokinetic and pharmacodynamic parameters

**DOI:** 10.1101/2020.04.24.059626

**Authors:** Aldenora Maria Ximenes Rodrigues, Rayla Kelly Magalhães Costa, Ranyelison Silva Machado, Stanley Juan Chavez Gutierrez, Francisco das Chagas Alves Lima, Aldeídia Pereira de Oliveira

## Abstract

The process involved in the research, discovery and development of drugs is characterized by high extensive and complex cost linked to scientific and technological innovations, and it is necessary to study and verify the progress of research carried out in the field that results in patent applications. *Aniba riparia (Nees) Mez* is a plant species often used for therapeutic purposes, where its pharmacological properties are associated to the presence of alkaloids called riparins. 5 synthetic analog compounds (riparins A, B, C, D, E and F) were developed from natural riparins. These molecules, natural and synthetic, showed several pharmacological activities in tests performed *in vitro and in vivo*, highlighting the Central Nervous System (CNS). The objective of this work was to evaluate the physical-chemical, pharmacokinetic parameters (absorption, distribution, metabolism, excretion and toxicity) and pharmacodynamic parameters (bioactivity and adverse reactions) of Riparin B by means of *in silico* computational prediction. *Online software* such as *Pre-ADMET, SwissADME, Molinspiration* and PASS *on line* were used for the analysis. Riparin B fits the characteristics of *druglikeness*, pharmacokinetic properties appropriate to the predicted patterns and activities within the scope for the treatment of AD, demonstrating a possible potential in the inhibition of AChE. Therefore, in silico results allow us to conclude that riparin B is predicted to be a potential future drug candidate, especially via oral administration, due to its relevant Drug-likeness profile, bioavailability, excellent liposolubility and adequate pharmacokinetics, including at the level of CNS, penetrating the blood-brain barrier.

## INTRODUCTION

The process involved in the research, discovery and development of drugs is characterized by high extensive and complex cost linked to scientific and technological innovations, and it is necessary to study and verify the progress of research carried out in the field that results in patent applications^[1]^.

Based on this principle, the pharmaceutical industry applies high investments in bioprospecting research, although it is aware that research on new drugs is a high-risk market. Thus, drug design strategies began to include molecular recognition studies in biological systems, assuming great importance, as they became fundamental bases for the understanding of properties such as potency, affinity and selectivity and structure-activity. And thus, the biotechnological tools associated with medicinal chemistry methods have gained a prominent role in the development of new molecules with biological activity^[2]^.

In order to avoid this failure, a set of ADME/Tox *in silico* filters was implemented in most pharmaceutical companies, aiming to discard substances, in the discovery phase, that are likely to fail later. This strategy tends to reduce the probability of failure, reducing time and resources used in research. And so, several softwares were developed that perform different analysis of molecules, inferring on the physicochemical, pharmacokinetic and pharmacodynamic parameters in the development stage^[3,4]^.

The species *Aniba riparia* is commonly used for medicinal purposes due to its pharmacological properties, attributed to alkaloids called riparins. Besides the natural molecules of this species, there are synthetic analogues known as riparins A, B, C, D and F, which share structures similar to natural molecules^[5]^.

Synthetic riparin analogues have already demonstrated antioxidant, including analysis of isolated mitochondria of the brain of mice^[6,7]^, antimicrobial^[8]^, antiparasitic^[9]^, as well as leishmanicide^[10]^, vasodilator^[7]^ and anti-inflammatory^[11]^ activities. Among the properties of synthetic riparins, their action in the Central Nervous System (CNS), such anxiolytic effects^[12]^ in anxiety models, without affecting the locomotion of the animals used in the experiment^[7]^, which make them target molecules of new studies aimed at obtaining therapeutic alternatives, and can be used in the treatment of AD.

However, there is a significant problem that still remains in the drug discovery procedure, particularly in the later stages of conducting research: the analysis of the properties of ADME (absorption, distribution, metabolism and excretion) and the evident toxicity of drug candidates. Over 50% of drug candidates fail due to poor analysis of ADME/Tox during a drug design^[2]^.

To evaluate the physicochemical, pharmacokinetic (absorption, distribution, metabolism, excretion and toxicity) and pharmacodynamic (bioactivity and adverse reactions) parameters of the Riparin B by means of in silico computational prediction.

## MATERIALS AND METHODS

This is an experimental, quali-quantitative research, aiming at the drug design by in silico analysis.

The design of the riparin B molecule was made through the *GaussView* 6.0 *software*, the structural parameters calculated through the *Gaussian* 09W program and transformed into MDL Molfiles by *Chem3D*, for use in the *software.*

The computational prediction for riparin B was performed through the *online software*: *Pre-ADMET®* (https://preadmet.bmdrc.kr/), *SwissADME®* (https://swissadme.ch), *Molinspiration® (*https://www.molinspiration).com/), in order to obtain relative results of physicochemical parameters (lipophilicity (logP), molecular weight, polar surface area, number of hydrogen bond donors and acceptors, number of rotary bonds and solubility in water), drug-likeness profile, pharmacokinetic profile (absorption, distribution, metabolism, excretion and toxicity) of the molecule; and *PASS on line (*http://www.pharmaexpert.ru/passonline/), which allows to predict results regarding the bioactivity profile (pharmacodynamics) and adverse reactions of the molecule.

The ADMET profiles of Riparina B were analyzed in comparison with the properties range for 95% of known drugs and the values were calculated by the the referred software server.

## RESULTS

Fig 1 shows the structural formula of the riparin B.

**Fig 1.**
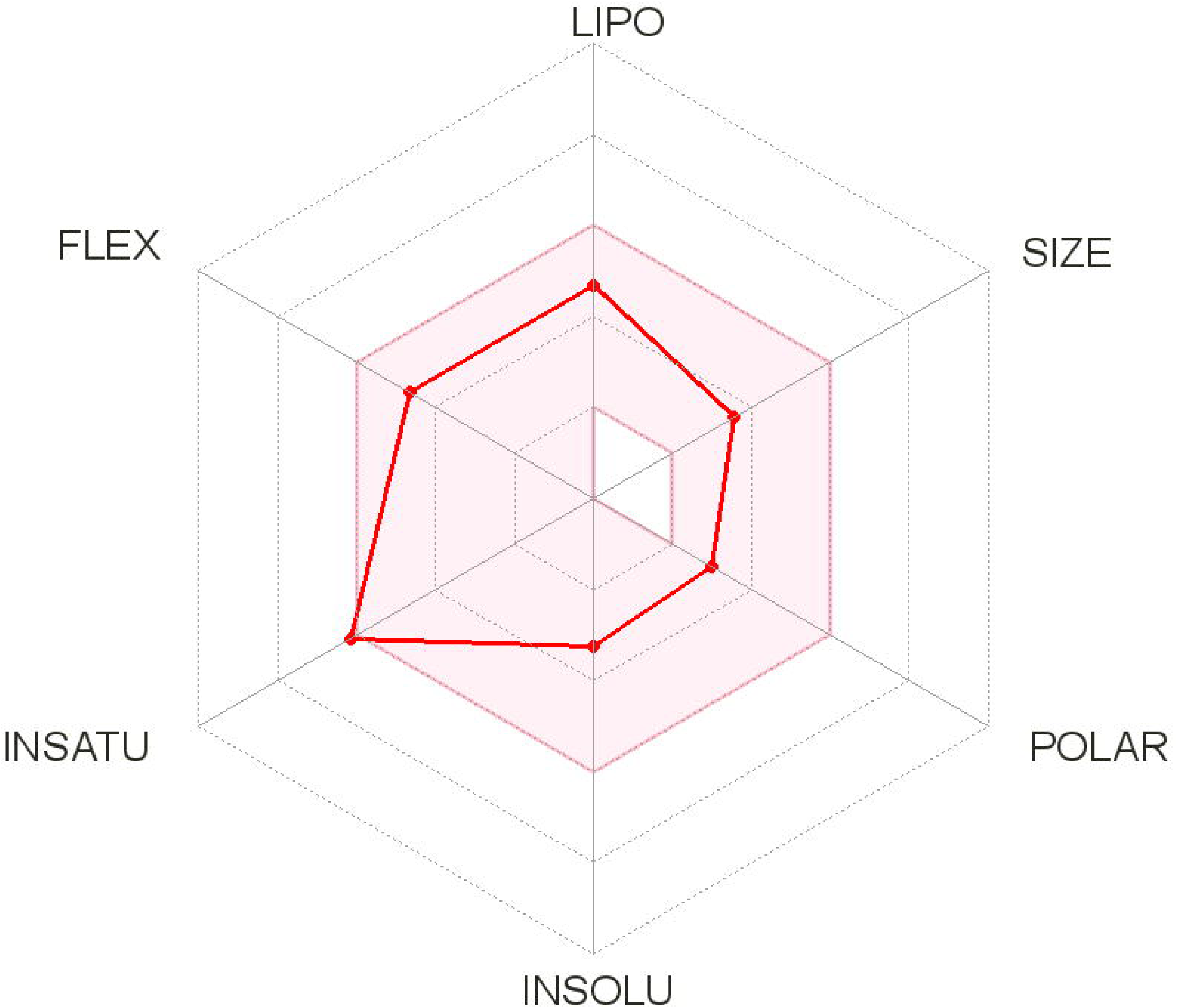
Structural formula of the riparin B.

Table 1 presents the physicochemical parameters, lipophilicity and solubility of riparin B predicted by Molinspiration, preADMET and SwissADME *software*.

**Table 1.** Physicochemical parameters, lipophilicity and solubility of riparin B.

| Physicochemical properties |  |
| --- | --- |
| Formula | C<sub>17</sub>H<sub>19</sub>NO<sub>3</sub> |
| Molecular weight | 285,34 g/mol |
| Number of atoms | 21 |
| Number of Arom. atoms. | 12 |
| Fraction Csp<sup>3</sup> | 0,24 |
| No. of rotating bonds | 7 |
| No. H bond Acceptors | 3 |
| No. of H bond Donors | 1 |
| No. of O+N (HBA) bonds | 4 |
| No. of OH + NH (HBD) bonds | 1 |
| Refractivity | 81,72 |
| TPSA (Topological polar surface area) | 47,56 Å<sup>2</sup> |
| Lipophilicity |  |
| P / w log (*i*LOGP) | 2,54 |
| P / w log (*X*LOGP3) | 2,68 |
| P / w log (*W*LOGP) | 2,68 |
| P / w log (*M*LOGP) | 2,56 |
| Log P / w (*SILICOS-IT*) | 3,41 |
| Overall average of the 5 predictions | 2,77 |
| Solubility in water |  |
| Log S (*ESOL*) | -3,26 |
| Solubility | 1,57e-01 mg/mL; 5,52e-04 mol/L |
|  | Soluble |
| Log S (*Ali*) | -3,33 |
| Solubility | 1,33e-01 mg/mL; 4,67e-04 mol/L |
|  | Soluble |
| Log S (*SILICOS-IT*) | -5,97 |
| Solubility | 3,02e-04 mg/mL; 1,06e-06 mol/L |
|  | Moderately soluble |

Legend: HBD - *Hydrogen Bond Donor*, HBA - *Hydrogen Bond Acceptor*.

Fig 2 is a diagram predicted by the *SwissADME software* which corresponds to the appropriate profile of the oral bioavailability of a drug.

**Figure 2.**
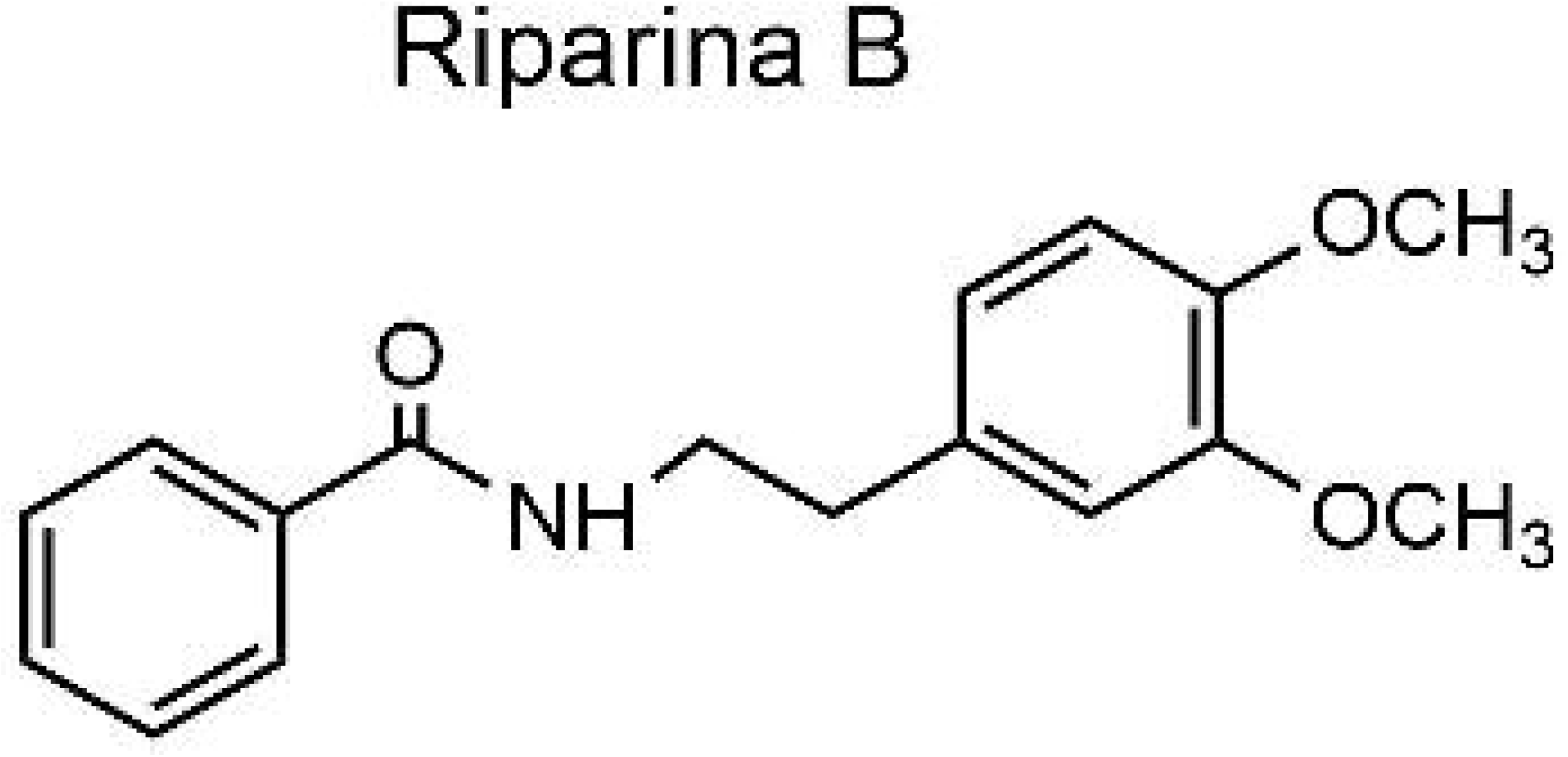
Oral bioavailability diagram (radar). Legend: LIPO - lipophilicity; SIZE - size; POLAR - polarity; INSOLU -insolubility; INSATU - unsaturations; FLEX - flexibity. The colored zone is the appropriate physical-chemical space for oral bioavailability.

Table 2 shows the *drug-likeness* profile predictions provided by the preADMET and SwissADME software.

**Table 2.** *Drug-likeness* profile.

| Similarity to drugs |  |
| --- | --- |
| *Lipinski* | Yes; 0 violation |
| *Ghose* | Yes; 0 violation |
| *Veber* | Yes |
| *Egan* | Yes |
| *Muegge* | Yes |
| *Lead rule* | Yes; 0 violation |
| MDDR | No; 1 violation |
| WDI Rule | Yes; 0 violation |
| Bioavailability score | 85% |
| **Medical Chemistry** |  |
| PANS | 0 alert |
| Brenk | 0 alert |
| Synthetic accessibility | 1,86 |

Legend: MDDR - MDL *Drug Data Report*; WDI - *World Drug Index*; PAINS - *Pan Assay* *Interference Structures*.

Table 3 shows the results related to the pharmacokinetic profile analyzed by the preADMET and SwissADME software.

**Table 3.**
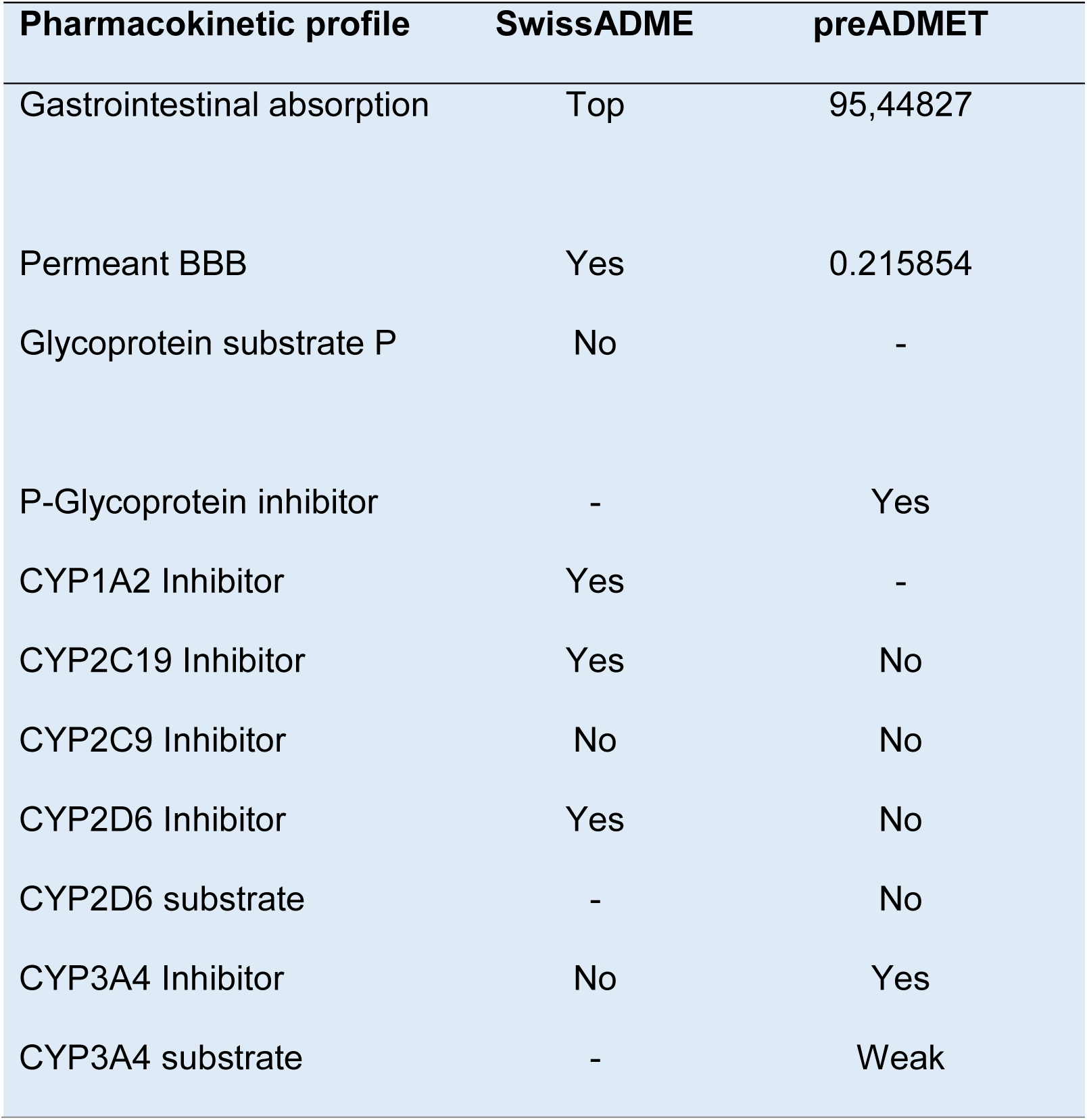
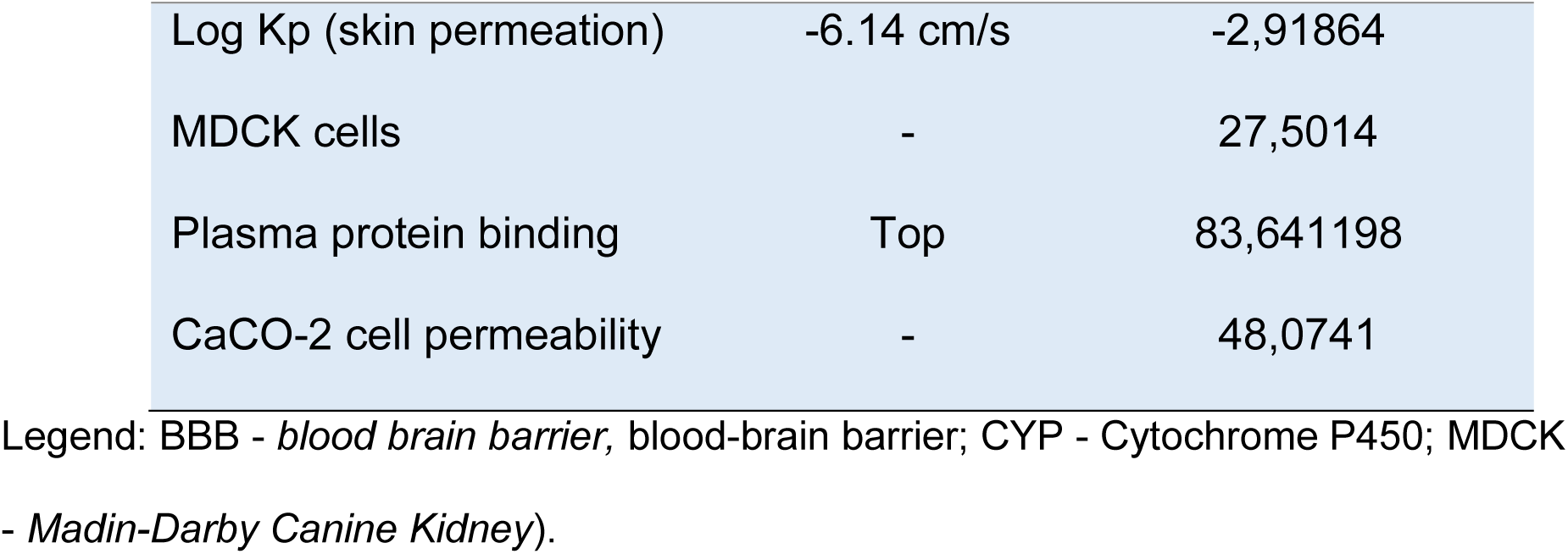
Pharmacokinetic profile of riparin B.

In table 4 there is a brief prediction of toxicity of the molecule by preADMET.

**Table 4.** Toxicity prediction of riparin B by preADMET.

| Toxicity |  |
| --- | --- |
| Ames Test | Mutagenic |
| *S. typhimurium* TA100 | Positive |
| *S. typhimurium* TA135 | Positive |
| Carcinogenic in mice | Positive |
| Carcinogenic in rats | Negative |
| Inhibition of hERG | Medium risk |

Legend: hERG - *human Ether-à-go-go-Related Gene*

Table 5 has information regarding the bioactivity *score* provided by the *Molinspiration software*.

**Table 5.** Bioactivity *Score* of riparin B. Legend: GPCR - Protein Coupled Receivers G.

|  |  |
| --- | --- |
| Binding to GPCR | -0.03 |
| Ionic channel modulation | -0.14 |
| Protein kinase/kinase inhibition | -0.13 |
| Connection to nuclear receivers | -0.29 |
| Protease inhibition | -0.12 |
| Enzyme Inhibition | -0.09 |

Table 6 shows the results of the probable pharmacological activities of riparin B in the CNS, predicted by the *software* PASS *on line.*

**Table 6.** Predicted pharmacological activities of riparin B.

| **Pa** | **Pi** | **Pharmacological activity** |
| --- | --- | --- |
| 0,876 | 0,010 | Treatment of phobic disorders |
| 0,456 | 0,005 | Dopamine release stimulant |
| 0,443 | 0,049 | 5-hydroxytryptamine release inhibitor |
| 0,341 | 0,058 | anti-amyloidogenic |
| 0,281 | 0,006 | Treatment of Huntington's disease |
| 0,299 | 0,068 | Antineurogenic pain |
| 0,333 | 0,105 | Treatment of dementia |
| 0,268 | 0,082 | Treatment of opiate dependence |
| 0,184 | 0,005 | Acetylcholine release stimulant |
| 0,185 | 0,019 | D2S dopamine antagonist |
| 0,188 | 0,023 | MAO Inhibitor |
| 0,275 | 0,116 | Acetylcholinesterase inhibitor |
| 0,279 | 0,132 | N-acetylneuraminate 7-O(or 9-O)-acetyltransferase inhibitor |
| 0,158 | 0,041 | Neuropeptide agonist |
| 0,156 | 0,049 | Acetyl-CoA C-acyltransferase Inhibitor |
| 0,297 | 0,212 | Neurotransmitter uptake inhibitor |
| 0,113 | 0,053 | Beta amyloid aggregation inhibitor |
| 0,078 | 0,025 | Butilcolinesterase Inhibitor |
| 0,119 | 0,067 | Neuropsin inhibitor |
| 0,090 | 0,049 | Adrenergics |
| 0,137 | 0,097 | Beta-amyloid protein antagonist |
| 0,196 | 0,168 | Antiparkinsonian |

Key: Pa – Probability to be active; Pi – probability to be inactive.

While table 7 predicts the possible and best targets of riparin B provided by *SwissADME software.*

**Table 7.** Prediction of riparin B targets, according to SwissADME.

| Target | Common Name | Target Class |
| --- | --- | --- |
| Acetylcholinesterase | AChE | Hydrolase |
| Delta opioid receptor | OPRD1 | Family A G-protein coupled receptors |
| Dopamine D1, D2 and D3 receptor | DRD1, DRD2 and DRD3 | Family A G-protein coupled receptors |
| Kappa opioid receptor | OPRK1 | Family A G-protein coupled receptors |
| GRM4 glutamate metabotropic receptor 1 and 4 | GRM1 and GRM4 | Family C G-protein coupled receptors |
| Monoamine oxidase A | MAOA | Family C G-protein coupled receptors |
| Serotonin 2a (5-HT2a), 2b (5-HT2b) and 2c (5-HT2c) receptor | HTR2A, HTR2B and HTR2C | Family A G-protein coupled receptors |
| Transient receptor potential cation channel subfamily A member 1 | TRPA1 | Voltage Dependent Ion Channel |
| Vanilloid receptor | TRPV1 | Voltage Dependent Ion Channel |

Table 8 presents the results of possible adverse reactions of riparin B also predicted by the PASS online software.

**Table 8.** Main expected adverse reactions of riparin B.

| Pa | Pi | Adverse reactions |
| --- | --- | --- |
| 0,908 | 0,004 | Galactorrhea |
| 0,861 | 0,008 | Orthostatic postural hypotension |
| 0,771 | 0,004 | Palpitation |
| 0,773 | 0,007 | Hypercholesterolemia |
| 0,781 | 0,045 | Tremors |
| 0,750 | 0,017 | Hyperglycemia |
| 0,746 | 0,018 | Hypotonia |
| 0,704 | 0,024 | Gastrointestinal bleeding |
| 0,711 | 0,037 | Atrial fibrillation |

Key: Pa – Probability to be active; Pi – Probability to be inactive.

## DISCUSSION / CONCLUSION

The use of in-silico models has been recognized in recent decades as being of fundamental importance in the area of research and development of drugs (R&D), due to its applications both in the evaluation of bioactive substances and in relation to their physicochemical and pharmacokinetic properties, giving rise to a new model of drug design with greater effectiveness and efficiency^[13]^.

Several pharmaceutical companies in different countries conduct their research related to ADME/Tox (Absorption, Distribution, Metabolism, Excretion/Toxicity) more efficiently through the introduction of combinatorial chemical technology, which allows the synthesis and screening of numerous compounds in the same time interval. This is only possible due to virtual screening (prediction systems), built by different programming languages forming databases with already known experimental data, allowing the screening of drug candidates, using the very drugs that have been approved and are already on the market^[14]^.

With the use of ADME/Tox prediction systems, it became easier to predict the action and behavior of numerous bioactive molecules that could become a drug, as a result this led to a better screening process of substances for research involving more advanced experimentation, reducing costs and time in drug development. In addition, in these screening systems it is possible to predict numerous failures that arise in the development of new drugs, sometimes foreseen in more advanced stages, such as clinics^[15,16]^.

Combinatorial chemistry has become gold in the pharmaceutical industry, since bioassays are guided by structural evaluation, observing structural similarities and/or interaction between new compounds, from the English *New Chemical Entity* (NCE) and receptors, as such revolutionizing the search for bioactive ingredients^[17,18]^.

Among all the parameters evaluated in this article, the discussion begins with explanations of the physicochemical properties of riparin B (fig. 1), presented in table 1. The search for understanding the physicochemical properties of small molecules has greatly increased. The understanding of these properties is necessary in the design of new pharmacological compounds with the ability to bind to various biological targets and present beneficial effects to the body, leading to the discovery of new treatments for diseases of more complex origin, such as Alzheimer’s disease^[19]^.

Some properties such as electronic distribution, size of the molecule, hydrophobicity, binding characteristics, presence of groups responsible for the biological activity of the molecule and flexibility are major influencers and with the ability to modulate the behavior of the molecule in a biological organism, including transport properties, bioavailability, affinity for proteins, metabolic stability, toxicity, among others^[20]^.

One of the physicochemical properties essential in the search for new drugs is the molecular weight (PM), which can be a great differential in relation to intracellular processes, such as intestinal absorption, penetration in the blood-brain barrier (BHE), elimination rate and interaction with molecular targets^[21]^. The analyzed molecule, riparin B, showed molecular weight with acceptable variability by all in silico filters provided by the *software.*

Another physicochemical characteristic of great importance obtained was with respect to the acid-base character of the molecule, determined by the ability to accept and donate protons H+^[22]^. Lipinski *et al.* ^[23]^ inferred that molecules that exhibit a lower number of hydrogen bond donor atoms - sum of *hydrogen bond donor atoms* O-H and N-H (HBD) and a higher number of hydrogen bond *acceptor atoms* - sum of *hydrogen bond acceptor atoms* O and N (HBA) have the most favorable ADME/Tox profile.

Ribparin B has shown to be an aceptor molecule, which infers that it has basic properties. This inference is of great relevance in relation to the pharmacokinetic process of absorption, more specifically possible absorption sites, since it is known that the main biological compartments have defined pH.

Among all the physicochemical properties of a micromolecule, the main ones capable of changing the pharmacotherapeutic profile are the ionization coefficient, expressed by pKa, which consists of the relative contribution of neutral and ionized species, already discussed, and the partition coefficient, expressed by the relation between hydro and liposolubility profile^[22]^.

Regarding the hydro and liposolubility profile of riparin B, it is possible to deduce that it is a liposoluble molecule with moderate water solubility (amphiphilic class), because the results obtained are within the variability accepted by different computational methods used.

Liposolubility (LogP) is a property of great significance and is used as an indicator of the oral bioavailability of drug candidate molecules, also constituting one of the main parameters of ADME/Tox^[24]^. In general, the optimization of the gastrointestinal absorption profile, through passive diffusion, after oral administration of a prototype candidate drug is achieved through the balance of its permeability and water solubility profile, known as Log P or Log D^[25]^.

Ribparin B presented an average of 2.77 for Log P, classified as optimal for good intestinal absorption, due to the balance between water solubility and the permeability rate by passive diffusion. Extreme values result in unbalance in these profiles, with capacity to negatively impact the oral bioavailability profile. In addition, the increase in lipophilicity values is involved in toxic properties such as blocking of CYP450 and hERG, as well as phospholipidosis induction^[24,26,27]^.

Thus, there is relevant evidence suggesting that controlling lipophilicity, among all the physical-chemical properties, within a defined ideal range, improves the quality of a molecule and, consequently, the probability of therapeutic success.

Besides LogP, the *software PreADMET* and SwissADME make available the TPSA of the molecule, often associated with the bonds that the structure is capable of making and which is also involved with modifications in oral permeability. This parameter is also used in association with the counting of rotational bonds and allows the analysis of the molecular flexibility, acting on the Drug-likeness profile of the molecule^[28]^.

Considering that the great majority of the active drugs by oral route are passively absorbed, having transpose the lipidic layer that constitutes the hydrophobic environment of the biological membranes, fig. 2 highlights important physicochemical properties necessary for the drug to reach plasma concentrations capable of reproducing the biological effect evidenced in *in vitro* and *in vivo* experiments. In the diagram it is possible to see that the characteristics of riparin B occupy only the colored zone, which is the appropriate physicochemical space for oral bioavailability.

In this study it was also possible to evaluate the Drug-likeness profile (table 2) of riparin B, through the physicochemical parameters of the molecule, such as PM, TPSA, HBA, HBD, Log P, number of atoms *in* general and aromatics atoms, fraction Csp3, number of rotative bonds and refractivity, in order *to* verify the similarity of riparin B with the other drugs already recognized and that are found in different in silico databases.

The best known rule that relates chemical structures to their biological activities is *Lipinski’s rule of five* or “rule of five”. It was developed to direct the choice of new drug candidate molecules and was also the pioneer in applying these rules to the drug-likeness profile of a given molecule with its physicochemical properties. According to this rule, for a given molecule to be permeable to cell membranes and also to have easy absorption by means of passive diffusion in the intestinal region, it needs to match the following parameters: LogP≤5; PM≤500; HBA≤ 10 and HBD≤5^[19,23]^. And riparin B meets all these requirements, without violation (Log P: 2.77; PM: 285.34; HBA: 4; HBD: 1).

In silico screening software also provides other filters to prove the *Drug-likeness* profile. Among them, the *Ghose* filter (160≤PM≤480; −0,4≤WLogP≤5,6; 40≤refrativity≤130; 20≤n° of atomos≤70), *Veber* (number of rotative bonds≤10; TPSA≤140), *Egan* (WLogP≤5,88; TPSA≤131,6), *Muegge (*200≤PM≤600; - 2≤XLogP≤5; TPSA≤150; number of aromatic rings ≤7; number of heteroatoms>1; number of rotative bonds≤15; HBA≤ 10 and HBD≤5), *Lead (*250≤PM≤350; XLogP≤3,5; number of rotative bonds≤7) ^[29]^ and displayed a qualified profile for a drug candidate.

There are also two Drug-likeness profile filters, (MDDR and WDI), made available by in-silico analysis from licensed databases that assign biological activities to drug-like compounds. The MDL Drug Data Report (MDDR), compiled from the patent literature, is a popular example. It contains several hundred distinct activities, some of which are therapeutic areas^[31,32]^. Concerning the MDDR filter, riparin violates a requirement, due to the presence of aromatic rings, and is considered as a possible medium profile drug. meanwhile for the WDI Rule, riparin does not violate rules and is similar to the drugs that belong to this database.

Drug-likeness is defined as a complex balance between various molecular properties and structure characteristics that determine whether a given molecule is similar to an oral medication with regards to bioavailability. These properties, especially hydrophobicity, electronic distribution, hydrogen binding characteristics, size and flexibility of the molecules and the presence of various pharmacophoric characteristics influence the behavior of the molecule in a living organism, including bioavailability, transport properties, affinity with proteins, reactivity, toxicity, metabolism, stability. Finally, it interferes with the efficacy relative to the pharmacokinetic profile of a molecule ^[33]^.

Responding to a need demonstrated by scientists to predict permeability and bioavailability properties, Martin^[34]^ has constructed a bioavailability score that seeks to predict the probability of a molecule having at least 10% oral bioavailability in rats or having measurable permeability in Caco-2 cells of 85% if the polar surface area (TPSA) is ≤ 75Å^2^; 56% if 75 <TPSA <150Å^2^ and 11% if TPSA is ≥150Å^2^. Riparin B showed an 85% probability due to TPSA value (47.56 Å^2^).

The objective of the results of medicinal chemistry is to support the daily efforts in drug discovery. Thus, the *SwissADME software* presents two complementary filters (PAINS and Brenk) for pattern recognition that allow the identification of potentially problematic fragments in the studied molecules. If there is any type of mentioned fragment found in the molecule under evaluation, the software indicates with alerts^[29]^. Taking these criteria into consideration, riparin B does not have this type of fragment, since there was no alert type to be considered.

Pan-assay interference compounds (PAINS) are chemical fragments that generally give false-positive results, as they tend to react unspecifically with numerous biological targets, rather than specifically connecting to a desired target^[35]^. The structural alert indicated by *Brenk* is purely based on the knowledge of a compilation of chemical parts known to be toxic, chemically reactive, metabolically unstable or with properties responsible for poor pharmacokinetics^[36]^.

Regarding the value of synthetic accessibility, riparin B presented a score of 1.86, shown to be an easily synthesized molecule. This value is a score based on the fragmented analysis of structures of more than 13 million compounds with the hypothesis that the more a molecular fragment is frequent, the easier it is to obtain the molecule. The score is defined between 1 (easy synthesis) and 10 (very difficult synthesis) ^[29]^.

Regarding the pharmacokinetic parameters (table 3), we obtained data on absorption (gastrointestinal (HIA), skin permeation (Log Kp), model of Caco-2 cells and MDCK cells), distribution (penetration of the blood-brain barrier and binding to plasma proteins), metabolism/biotransformation/excretion (substrate and cytochrome P450 inhibitor).

Among the numerous in vitro methods used in the drug selection process to evaluate the intestinal absorption of drug candidates, the Caco-2 (human adenocarcinoma colorectal cell culture) and MDCK (*Madin-Darby Canine Kidney*) cell models have been recommended as reliable for predicting the oral absorption of drugs. In addition, the HIA (*Human Intestinal Absorption)* in silico model and the skin permeability model can predict and identify potential drugs for oral and transdermal administration^[37]^.

Regarding these parameters, riparin B demonstrated high gastrointestinal absorption potential (GIA), which corroborates the basicity of the molecule and relevant skin permeation values, both by *preADMET* (−6.14 cm/s) and *SwissADME* (−2.91864) *software.*

Oral administered drugs are preferably developed, due to market, convenience and safety. After oral administration, the drug goes through different processes, among them: it is dissolved and solubilized in the gastrointestinal tract so that it can be absorbed in the stomach or through the intestine. The latter, called human intestinal absorption is one of the most important for drugs that act through this route^[38]^. Thus, we can predict that riparin B has a significant potential to become an oral drug.

The prediction of the permeability coefficient (Kp) for the absorption of molecules by the epidermis of mammals is based on the linear model built by Potts and Guy^[39]^ Thus, the more negative the log Kp, the less the molecule permeates, for a example, diclofenac, good topical anti-inflammatory with a log Kp (−4.96 cm/s).

The results predicted by the Coco-2 (∼48.0) and MDCK (∼27.5) cell models also pointed out that riparin B fits as an average permeability molecule (20 ∼ 70 %), according to the category proposed by Yamashita and collaborators^[40]^.

Still regarding absorption, one of the results found in this study was that riparin B may act as a P-glycoprotein (P-gp) inhibitor and not as a substrate. This result fundamental to deduce about active efflux through biological membranes, since P-glycoprotein constitutes a class of efflux or secretion transporters that act as a barrier to absorption in numerous compartments, such as in the gastrointestinal membranes and lumen wall or in the membranes of the brain. An important role of P-gp is to protect the CNS from xenobiotics and if the molecule can act as an inhibitor, it reinforces the possibility of crossing the blood-brain barrier^[41]^.

Regarding distribution, the penetration of the blood-brain barrier can provide information on the therapeutic potential of the drug in the CNS and the model of binding to plasma proteins provides data on an effective distribution. According to the predicted results, riparin B has the ability to overcome the blood-brain barrier (∼0.21) and can be characterized as a substance with medium absorption (0.1-2.0) and binding power to plasma proteins (∼83%), which can be considered a weakly bound chemical substance (less than 90%).

Predicting penetration into the blood-brain barrier means predicting that the molecule is able to pass through this barrier is crucial in the pharmaceutical sphere and in this study, since its main focus is the treatment of Alzheimer’s disease^[42]^.

The part of the drug that becomes available for diffusion through the membranes, and for pharmacological interaction in the body is that which is not bound to plasma proteins, thus a high affinity for plasma proteins directly influences the pharmacological activity and biodistribution of this drug^[37]^.

Knowledge about the interaction of molecules with cytochrome P450 (CYP) is also essential. This superfamily of isoenzymes participates in a fundamental way in the metabolism, biotransformation and elimination of drugs. It is estimated that 50 to 90% of therapeutic molecules can be substrates of the five major isoforms (CYP1A2, CYP2C19, CYP2C9, CYP2D6, CYP3A4) ^[43]^.

Thus, it was possible to predict that riparin B proved to be a weak substrate of only the CYP3A4 isoform. Regarding the inhibition of the isoforms, the results of the *software* were contradictory, as *preADMET* reported that riparin B inhibits only the CYP3A4 isoform, whereas *SwissADME* predicted that the molecule inhibits the CYP1A2, CYP2C19 and CYP2D6 isoforms.

Inhibition of these isoenzymes is one of the main causes of drug interactions related to pharmacokinetics and may cause drug interactions, toxic effects or adverse effects due to less purification and accumulation of the drug or its metabolites in the body^[43]^.

Toxicity is one of the final parameters of the ADME/Tox analysis of the molecule in question. Considering the in silica toxicity exhibited by the *preADMET software*, riparin B displayed the following results (table 4): positive for the two strains of *S. typhimurium* used in the Ames test, implying it as mutagenic, negative for carcinogenicity in rats and positive for carcinogenicity in mice and showed medium risk of hERG inhibition(cardiotoxicity).

The *Ames* test is a simple method that detects mutagenicity of a substance by making use of various strains of *Salmonella typhimurium* bacteria that carry mutations in genes involved with histidine synthesis, the variable tested by the *software* is the ability of the mutagen to cause a reversal of growth in a histidine free medium^[44]^.

Computational modeling is often used to select new molecules for therapeutic purposes, based on the most relevant biological properties for pharmacological interaction. *Molinspiration software* provides bioactivity scores of molecules with respect to different cell receptors, such as ionic channels, GPCRs, enzymes, proteases, kinases and nuclear receptors^[45,46]^.

There is a ranking to be followed, where there is the inference that active substances are those that *score* >0, moderately active substances *score in the* range of −5.0 to 0 and inactive compounds *score <-5*.0^[47, 48]^. Through table 5, it was possible to predict that the riparin B molecule presents as a moderately active substance. It is important to mention the bioactivity of enzyme inhibition, important in the treatment of Alzheimer’s disease, since AChE inhibitors constitute a class of treatment.

The results obtained through the PASS *online software* are based on the computational learning method, the English “*machine learning methods*”, which makes use of multilevel descriptors and Bayesian algorithm, with the ability to predict the activity and inactivity probabilities for more than 4000 biological activities from a complete analysis of the biologically active molecule structure-activity^[49,50]^.

Important predictions of the pharmacological potential of riparin B were observed in table 6, focusing on activities directly related to the CNS. To analyze the results, there is a score to be used: activity with higher probability of occurrence (Pa>0.7), probable probability of occurrence (0.5< Pa<0.7) and unlikely probability of occurrence (Pa<0.5). Thus, it is predicted which pharmacological activities are very likely, probable or unlikely for molecules previously tested in *in vivo* experiments^[51]^.

The activities demonstrated that are related to Alzheimer’s disease include: dementia treatment, acetylcholine release stimulant, anti-amyloidogenic, acetylcholinesterase inhibitor, butylcholinesterase inhibitor, Beta amyloid aggregation inhibitors and β-amyloid protein antagonist (Table 6).

In addition, the *SwissADME software* has listed possible targets for the riparin B molecule (table 7). Among them, it is worth mentioning the acetylcholinesterase, target of several drugs commonly used in the treatment of Alzheimer’s disease.

Like almost all drugs, the prediction for riparin B inferred possible adverse reactions (table 8), with galactorrhea and orthostatic postural hypotension among the most likely adverse reactions. These reactions are likely to occur due to failures in the biotransformation process, specifically in the inactivation of drugs for subsequent excretion.

The molecular modeling strategy addresses different predictive models in order to mimic and get as close as possible to the properties that influence the administration of drugs, especially orally. In this way, it is a useful tool in the primary design of bioactive molecules with optimized pharmacokinetic properties, this makes it very useful in the design of new drug candidates in research laboratories^[13]^.

Therefore, in silico results allow us to conclude that riparin B is predicted to be a potential future drug candidate, especially via oral administration, due to its relevant Drug-likeness profile, bioavailability, excellent liposolubility and adequate pharmacokinetics, including at the level of CNS, penetrating the blood-brain barrier. It is also assumed that it can become a possible drug for the treatment of Alzheimer’s disease, since it targets the enzyme Acetylcolinesterase.

